# The entropic heart: Tracking the psychedelic state via heart rate dynamics

**DOI:** 10.1101/2023.11.07.566008

**Authors:** Fernando E. Rosas, Pedro A.M. Mediano, Christopher Timmermann, Andrea I Luppi, Diego Candia-Rivera, Reza Abbasi-Asl, Adam Gazzaley, Morten L. Kringelbach, Suresh Muthukumaraswamy, Daniel Bor, Sarah Garfinkel, Robin L. Carhart-Harris

**Author notes:** Equal contribution (Fernando E. Rosas, Pedro A.M. Mediano).

## Abstract

A growing body of work shows that autonomic signals provide a privileged evidence-stream to capture various aspects of subjective and neural states. This work investigates the potential for autonomic markers to track the effects of psychedelics — potent psychoactive drugs with important scientific and clinical value. For this purpose, we introduce a novel Bayesian framework to estimate the entropy of heart rate dynamics under psychedelics. We also calculate Bayesian estimates of mean heart rate and heart rate variability, and investigate how these measures relate to subjective reports and neural effects. Results on datasets covering four drugs — lysergic acid diethylamide (LSD), dimethyltryptamine (DMT), psilocybin, and sub-anaesthetic doses of the dissociative agent ketamine — show consistent increases in mean heart rate, high-frequency heart rate variability, and heart rate entropy during the psychedelic experience. Moreover, these effects have predictive power over various dimensions of the psychedelic experience. Changes in heart rate entropy were found to be correlated with increases in brain entropy, while other autonomic markers were not. Overall, our results show that a cost-efficient autonomic measure has the potential to reveal surprising detail about subjective and brain states, opening up a range of new research avenues to explore in both basic and clinical neuroscience.

---

**C**onscious experience is supported by the confluence of numerous bodily processes, which involve not only the brain, but also other systems such as the autonomous nervous system. Indeed, there is growing evidence supporting a fundamental role of the autonomic system in the generation of subjective experience and the sense of selfhood (1–3). It is now becoming clear that autonomic function is not restricted to homeostatic regulation and ‘fight or flight’ responses, but is also critically involved in cognitive processes such as attention, memory, and decision-making (4, 5), as well as in experiential phenomena such as emotional processing and regulation (6, 7) and body awareness (8, 9). Thus, the autonomic system has the potential to provide valuable insights into the physiological substrates that support human cognition (10) — which is also attractive from a practical perspective, considering that reliable measurements of heart activity are relatively simple and inexpensive, compared to measurements of brain activity.

Building on the above evidence, here we investigate the potential for autonomic markers to track the effects of psychedelic substances — potent psychoactive drugs capable of inducing profound changes in perception, cognition, and conscious experience (11). Psychedelics are exceptional tools for inducing perturbations to human consciousness, which is critical for exploring the mechanisms linking mind and brain (12, 13). More practically, evidence from phase II clinical trials suggests that psychedelic drug administration combined with psychotherapy may have efficacy for treating various mental health presentations, including depression, end-of-life distress, tobacco addiction, and alcoholism (14, 15). It is noteworthy that the majority of the efforts (such as Refs. (16–21)) trying to explain how psychedelics work has been mainly focused on the brain, broadly neglecting the rest of the body. Considering how the psychedelic experience affects these cognitive and experiential processes, and how relevant these effects may be for their clinical efficacy, the autonomic nervous system offers an essential and complementary evidence stream to enrich our understanding of how psychedelics act to produce their therapeutic and consciousness-altering effects.

This work also takes inspiration from intriguing parallels that exist between empirical findings related to psychedelic brain dynamics and general investigations on heart rate variability (HRV) — a popular way of studying the autonomic nervous system by assessing how heart rate changes over time (22–24). On the one hand, research has found that while classic psychedelics (e.g. lysergic acid diethylamide [LSD], psilocybin, and dimethyltryptamine [DMT]) act primarily on the 5-HT_2A_ receptor, and the atypical psychedelic ketamine exerts its effects primarily via NMDA receptor antagonism (25), they both reliably disrupt important markers of spontaneous brain function at wakeful rest evidenced, e.g., by reductions of alpha power and large-scale network integrity (26–29). This, in turn, induces a mode of brain function characterised by an enhanced brain entropy (i.e. more diverse and less stereotyped patterns of activity), which supports the enriched experience typical of the psychedelic state as suggested by theoretical (16, 19, 30, 31) and empirical (32–34) work. Consistently, but in another or-gan — the heart — a vast body of work has established a link between the temporal variability of heart beats and health, where such variability is high in healthy controls and reduced in a wide range of conditions (23). These empirical findings support the view that the heart is not a mere metronome, but rather a flexible organ that dynamically supports interactions with a complex and changing environment (35). Interestingly, metrics of dynamical complexity have been found to extract valuable information from the non-linear aspects of heart rate fluctuations (36, 37). By contemplating these two lines of research, it is then natural to ask: is it possible to measure the variability of heart dynamics in a way that is analogous to how brain entropy is measured? If so, do psychedelics increase this ‘heart entropy’ too? Would changes between brain and heart entropy be correlated, and would they be predictive of subjective experience?

Here we address these questions by leveraging a novel Bayesian method to estimate the entropy of heart rate (HR) dynamics. Our method provides estimates of HR entropy with high temporal resolution, thus allowing us to effectively track psychedelic drug action on cardiac activity over time. By investigating multiple datasets covering four different drugs (the classic serotonergic psychedelics LSD, DMT, and psilocybin, as well as the dissociative agent ketamine at sub-anaesthetic dosage), our results reveal that psychedelics consistently increase heart rate entropy, and that this increase provides information for predicting subjective reports that is complementary to known brain-based biomarkers. These findings open new opportunities to use autonomic signals for tracking the psychedelic state, with important implications for future clinical applications.

## Results

### A. Bayesian estimation of heart rate dynamics under psychedelics

The first step in our analysis was to validate the proposed Bayesian method to estimate heart dynamics from electrocardiogram (ECG) time series data of subjects under the effects of psychedelic drugs. For this, we used four datasets pertaining to LSD, DMT, psilocybin, and ketamine in healthy human volunteers (N = 20, 12, 15, and 20, respectively). Of these, the LSD dataset consists of a stable period occurring approximately four hours after administration, while the other datasets consist of a continuous period including before and after drug injection, hence reflecting the pharmacokinetics of the drug’s action (see *Methods* for details).

For each dataset, we extracted inter-beat intervals and used this information to reconstruct heart rate dynamics with a novel Bayesian approach (see *Methods*). Briefly, our method describes heart dynamics by formulating a posterior distribution over sequences of heart rate values — henceforth called ‘heart rate trajectories’ — conditioned on the observed sequence of inter-beat intervals. We then efficiently sample this posterior distribution with a Markov chain Monte Carlo (MCMC) approach to obtain 500 heart rate trajectories for each subject on each condition (either drug or placebo), some of which are illustrated in Figure 1. As a comparison, we also extracted heart rate dynamics via the standard (frequentist) approach, which provides a single trajectory (red curve in Figure 1). When compared with the standard approach, it is observed that the proposed Bayesian method provides a both a smooth average reconstruction of the HR time-series, as well as an ensemble of trajectories that captures the overall variability. Also, as expected, the data from DMT, psilocybin and ketamine show large changes in the minutes following injection, while the data from LSD exhibit less pronounced changes, possibly due to the fact that ECG recordings in this condition took place at the plateau of subjective effects (i.e. after the peak).

**Fig. 1.**
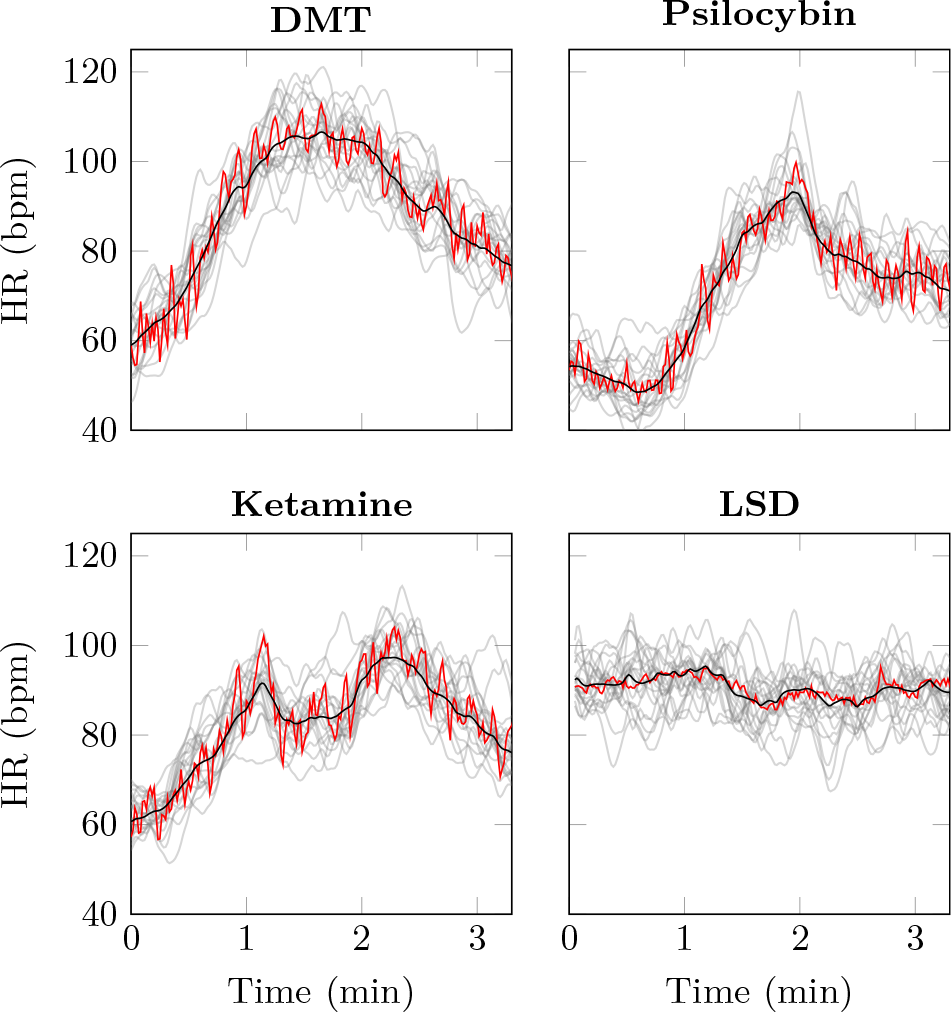
Illustration of our Bayesian method to estimate heart rate dynamics. Each panel shows the data of a single sample subject, for which we plot 40 heart rate trajectories sampled from the model’s posterior (gray), as well as their mean (black). The result of a standard frequentist estimation of the heart rate trajectory is also plotted for comparison (red). Injection of the drug takes place at *t* = 0 for the DMT, psilocybin, and ketamine data, while for LSD the data corresponds to 4 hours after injection.

### B. Consistent changes in autonomic markers between drug and placebo

After confirming the suitability of our approach, we used the ensemble of trajectories obtained in the previous step to build a new Bayesian estimator of the entropy of heart rate dynamics (see *Methods* for details). Crucially, following Bayesian principles, we compute average entropy as the average of the entropies of each of the 500 sampled HR trajectories (which, due to the non-linearity of entropy, is not equivalent to the entropy of the mean HR trajectory). Another innovation in our approach is to estimate entropy by training and testing the model on different datasets, which relies on the particular capabilities of the Context Tree Weighted (CTW) algorithm (38): while standard methods (e.g. the Lempel-Ziv algorithm (39)) estimate entropy by quantifying the diversity of observed patterns (40), the CTW algorithm allows us to assess pattern diversity in a signal using the patterns of another signal as reference. Accordingly, the CTW algorithm allowed us to estimate entropy by training over the data of the whole placebo session, and testing over small windows of data from either drug or placebo session.

Equipped with this method, we sought to investigate whether HR biomarkers could track the evolution of the psychedelic state. We calculated HR entropy over overlapping windows of 60 samples, corresponding to 60 seconds of data. Additionally, we calculated the mean heart rate, and the low- and high-frequency heart rate variability (HRV) (see *Methods*). The results were then analysed via cluster-statistics (41) looking for differences over the time course of either drug or placebo sessions with respect to their baseline for the DMT, psilocybin, and ketamine datasets, and paired t-tests for the LSD dataset (see *Methods*). Results show that HR markers exhibit significant changes after drug administration for all compounds, with increases in mean HR, high-frequency HRV, and HR entropy (Figure 2). Increases in low-frequency HRV are observed only in DMT and psilocybin. Increases in mean HR appear earlier for all drugs, followed by increases in HRV and HR entropy. Interestingly, results in the following sections illustrate how different autonomic markers have distinctive predictive power over neural and subjective effects.

**Fig. 2.**
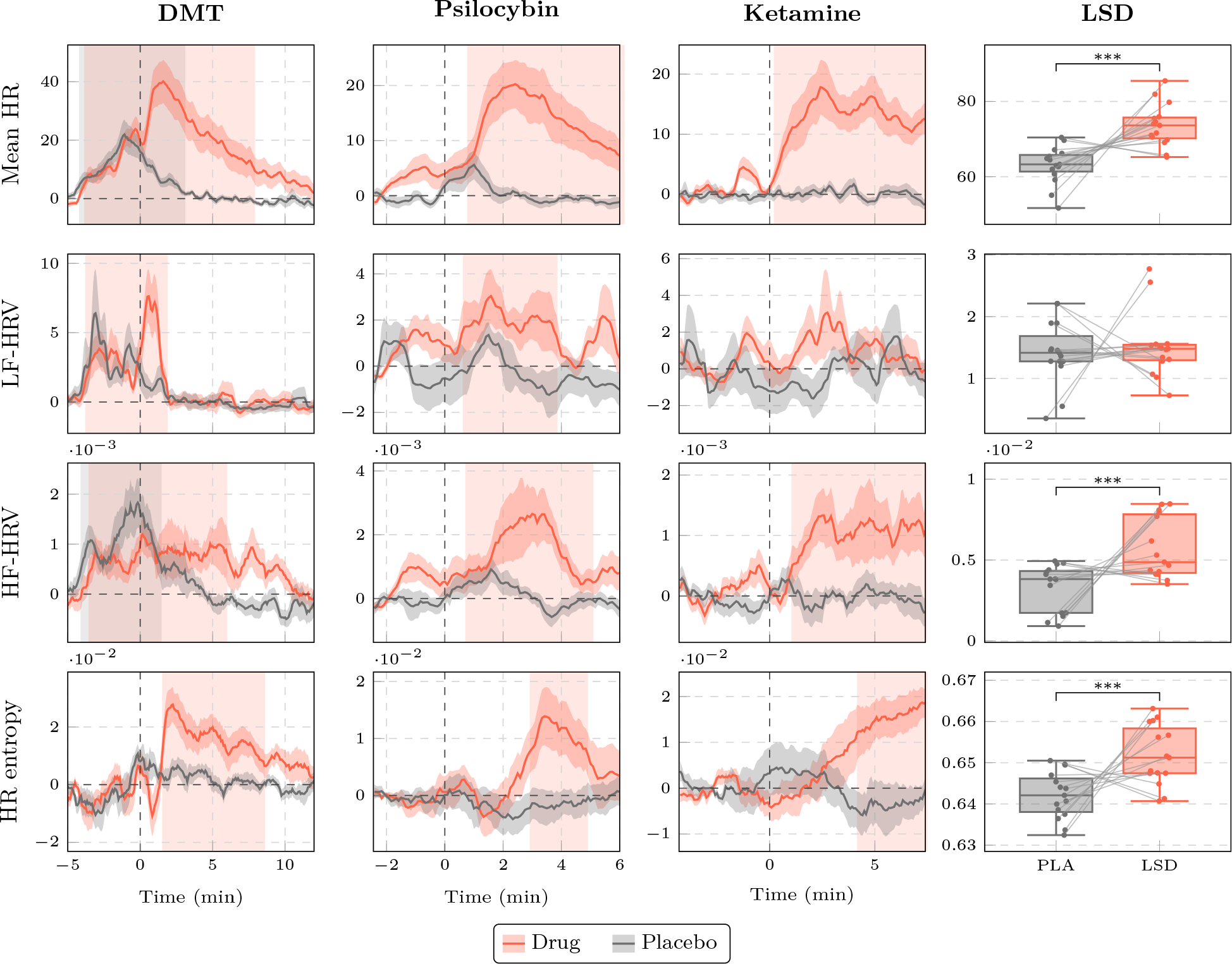
Heart rate biomarkers change under psychedelics. *Columns 1-3:* Profiles of temporal evolution of mean heart rate, low (LF-HRV) and high (HF-HRV) frequency heart-rate variability, and heart-rate entropy for DMT, psilocybin, and ketamine. Each biomarker is normalised by its average value before drug injection. Coloured windows correspond to statistically cluster-corrected significant differences (*p <* 0.05) of either drug (red) or placebo (gray) with respect to their baseline. *Column 4:* Changes of the same biomarkers for LSD (versus placebo) 4 hours after its injection (∗∗∗ : *p <* 0.001). Note that no pharmacokinetic profiles are available for this compound.

### C. Predictive power of autonomic markers evolves over time

After observing consistent changes in various markers of heart rate dynamics due to psychedelics, we sought to investigate how these effects are related to drug effects on the brain and on subjective experience as they unfold over time. As a proof of concept, we performed exploratory analyses on the predictive ability of these markers in the DMT dataset, motivated by the short-lived dynamics of DMT action (that are fully captured within the data) and the rich questionnaire data available from that study. Considering the exploratory nature of these analyses, we focus the report of our findings in the resulting effect sizes (in particular, explained variance *R*^2^), while providing indications of significance testing only for reference.

We first studied how much of the between-subjects variance of brain entropy changes could be explained by HR entropy. For this analysis, we calculated the brain entropy using the EEG data recorded during the DMT experience with the same CTW algorithm used on the heart rate (see *Methods*). We then calculated the proportion of explained variance (*R*^2^) of brain entropy from changes in HR entropy, at different timepoints during the DMT experience and with different lags between heart and brain signals. Results reveal two periods of substantial correlation between HR and brain entropy (Figure 3a): one during the peak experience, 0-5 minutes after injection, and another one 9-12 minutes after injection. In contrast, the mean heart-rate shows a much smaller *R*^2^, highlighting the superior predictive power of HR entropy metric over mere HR changes.

**Fig. 3.**
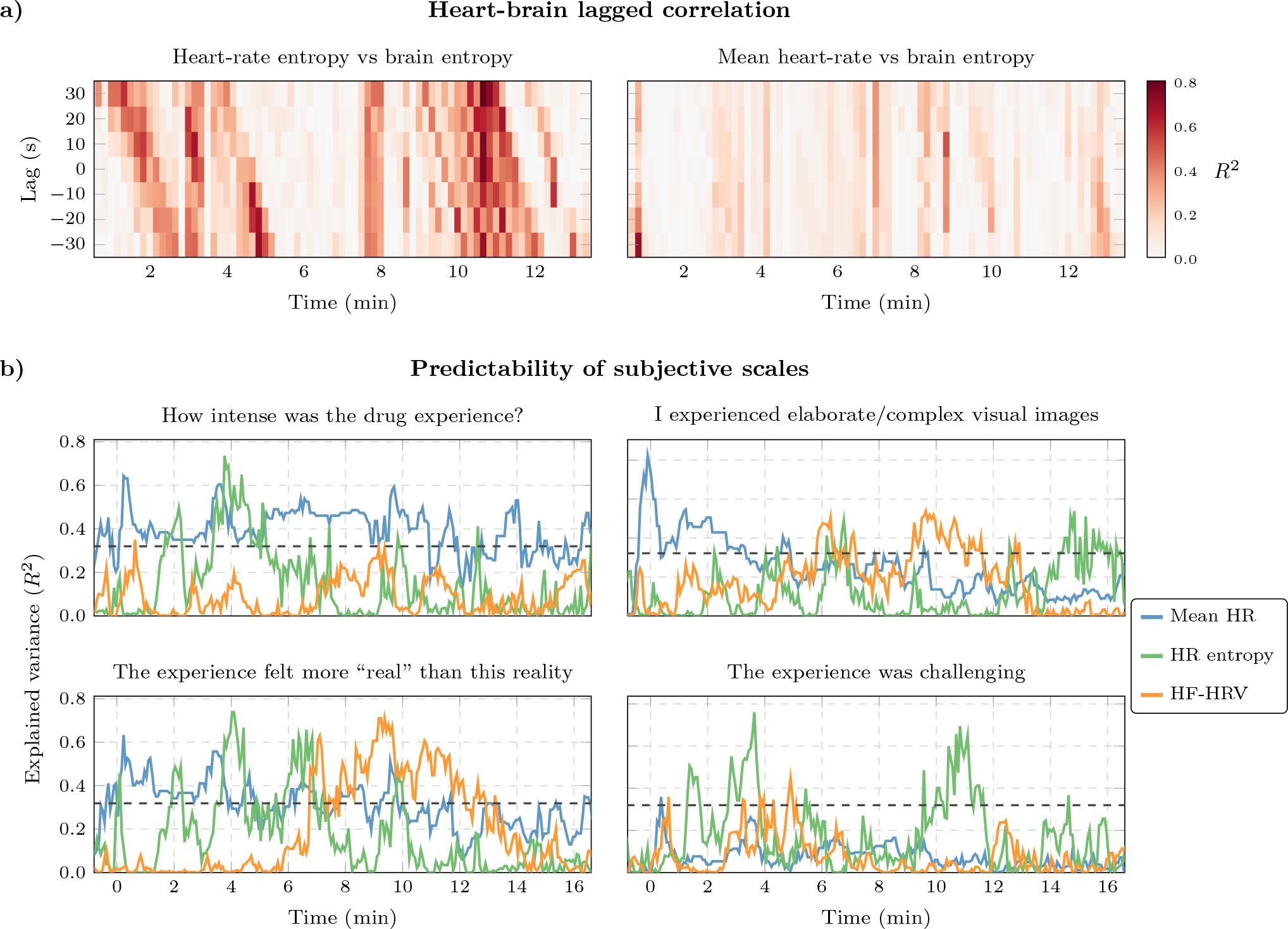
Predictive power of heart-rate biomarkers can change dynamically. **a)** The explained variance (*R*^2^) of changes in brain entropy predicted via changes in HR entropy reveals two periods of substantial correlation between them during the DMT experience. In contrast, changes in mean heart-rate are much less correlated with changes in brain entropy. Each block corresponds to a 10-second epoch, considering lags of up to 30 seconds with respect to HR entropy. Heatmaps thresholded by non-parametric confidence intervals can be found in Supplementary Figure 6. **b)** Autonomic markers can explain up to 70% of the between-subjects variance of various dimensions of the DMT experience, which were rated retrospectively via a single visual-analoge scale score. Those four subjective scales represent different types of relationships: intensity dominated by mean HR, challenging experience by HR entropy, complex imagery alternating between HRV, HRM, and entropy at different times. Dashed black line represents 95^th^ percentile of the null distribution (equivalent to the significance threshold of an uncorrected one-tailed hypothesis test with *α* = 0.05).

We then investigated the predictive power of autonomic makers over behavioural reports. For this, we used questionnaire data in the form of numerical values (from visual analog scales) that retrospectively characterise the whole psychedelic experience. Similarly, as with brain entropy, we calculated the explained variance within a rank regression using the score of each subject as dependent variable and the value of the change of heart rate marker as independent variable. Results show that different markers have distinctive predictive power at different timepoints, which can explain up to 70% of the variance between subjects in the reports of various dimensions of the DMT experience (Figure 3b).

### D. Topographical brain-heart association and complementarity in predictive power

Finally, we sought to investigate how heart entropy relates to changes in brain entropy in specific areas, and if these associations lead to a redundant or complementary ability to predict subjective responses. For this, we carried out exploratory analyses focused on the LSD dataset, which includes MEG data (thus, superior signal-to-noise ratio and spatial localisation than EEG) plus anatomical MRI data to perform source reconstruction, and the largest number of samples considering number of subjects (20), conditions (drug vs placebo), and settings (4) (see *Methods*), leading to 160 data samples — which can be exploited via linear mixed-effect modelling. Please note that the other datasets were not suitable for these analyses, as the DMT data is low-density EEG and the psilocybin and ketamine datasets do not have high-quality subjective data.

As a first analysis, we calculated the changes in brain entropy via the CTW algorithm similarly as in the previous section, but this time applied on source signals reconstructed at the centroids of the regions in the *automated anatomical labelling* (AAL-90) atlas (42) (see *Methods*). We then estimated the relationship between changes in heart and brain entropy via linear mixed-effects modelling using heart entropy as the dependent variable, brain entropy and condition (drug or placebo) as fixed effects, and non-nested random intercepts for subject ID and setting (see *Methods*). The significance of the association was measured via a log-likelihood test comparing this model against a similar model using only condition as fixed effect, and the resulting p-values were corrected via a Benjamini-Hochberg false discovery rate procedure (43). This analysis reveals that changes in heart and brain entropy are positively correlated in various brain areas located predominantly in regions of the default-mode network and sensorimotor networks (Figure 4a).

**Fig. 4.**
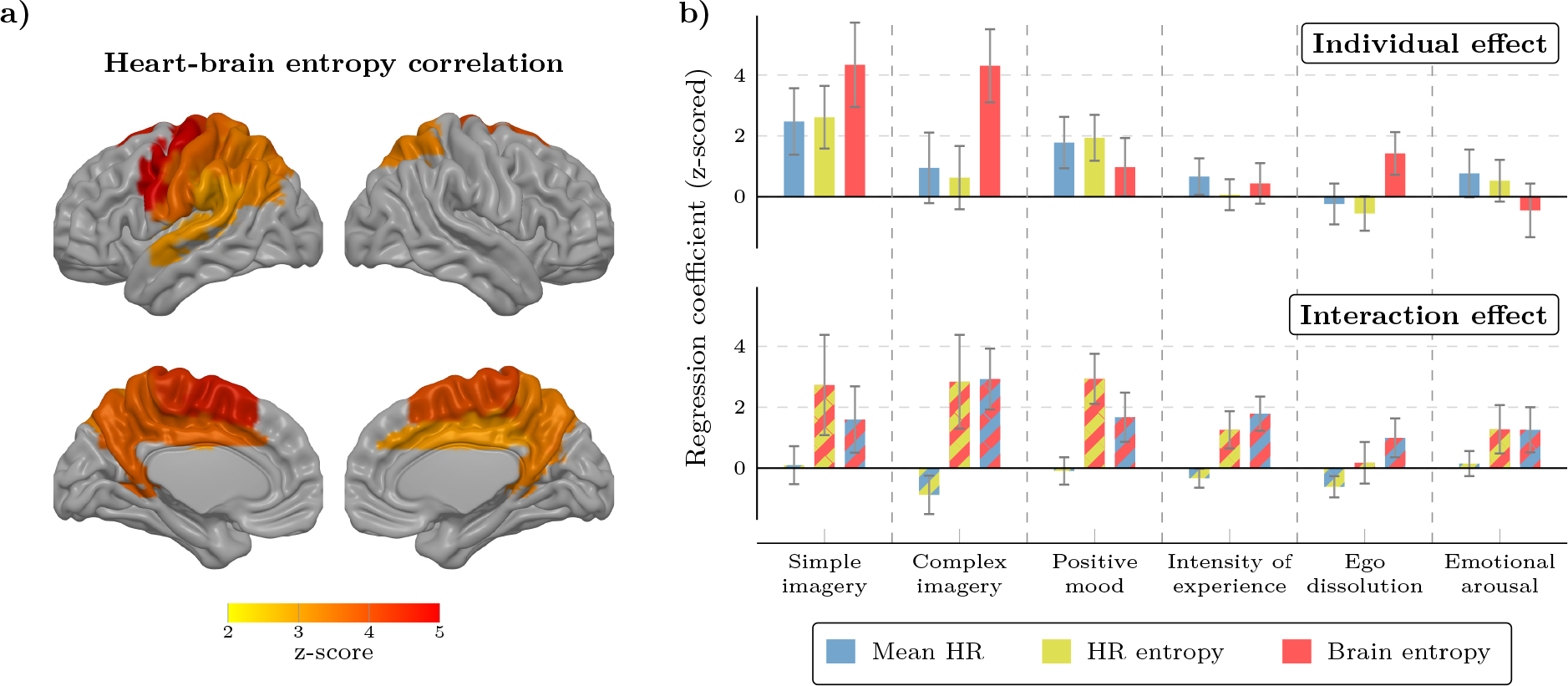
Relationship between heart and brain entropy changes under LSD. **a)** 17 regions from the automated anatomical labelling (AAL) parcellation exhibit significant statistical associations (after multiple comparison correction) with the observed changes in heart entropy. Areas include precuneus, mid cingulate, and sensorimotor regions. **b)** Predictive power of brain and autonomic markers over various aspects of the subjective experience after LSD administration, as shown via linear mixed-effect modelling. Top row corresponds to the drug-related effect of a single heart marker (i.e. interaction between the marker and the drug), and bottom row corresponds to the combined effect of two markers (i.e. three-way interaction between drug and two markers). Substantial positive interactions are observed between brain and autonomic markers. For comparability, all markers are z-scored before regression. Error bars represent the 95% confidence intervals of the coefficient estimates.

Next, we investigated the predictive power of heart and brain markers over subjective responses. For this purpose, we constructed linear mixed models to predict the scores on visual analog scales related to six features of the LSD experience: simple and complex imagery, positive mood, intensity of the experience, ego dissolution, and emotional arousal. Each model considered one of these subjective scores as the dependent variable, and used heart and brain biomarkers as predictors,^∗^ and random intercepts for subject ID and setting (see *Methods*). We consider two types of models: using a single marker as predictor, and using two markers and their interaction. When considering markers individually (Figure 4b, top), we found that brain entropy is the strongest predictor of simple and complex imagery and ego dissolution, HR entropy of positive mood, and mean HR of intensity and emotional arousal. When considering pairs of predictors (Figure 4b, bottom), results reveal various substantial positive interactions, particularly between brain entropy and the autonomic markers, which suggests that knowing the state of the autonomic system substantially increases the predictive power over subjective scores.

## Discussion

In this paper we put forward a new method to compute markers of heart-rate dynamics and showcased it on data from subjects experiencing the effects of different psychedelic compounds, which let us probe this altered state of consciousness from the lens of the autonomic nervous system. Results show that psychedelic drugs induce consistent increases in mean heart rate, high-frequency HRV, and HR entropy. This is a relatively uncommon pattern, as increases in HR are typically accompanied by decreases in heart variability (both spectral and/or entropic), while simultaneous increases of both have been observed in profiles associated to emotions such as ‘ joy’ (44) — which are hard to elicit in the laboratory.

These autonomic changes were found to have predictive power over various dimensions of the psychedelic experience. Additionally, changes in HR entropy were found to be correlated with changes in brain entropy, while having complementary predictive power on particular subjective responses. Overall, these findings illustrate how autonomic signals, when assessed with adequate analytical tools, can provide valuable information on various aspects of the psychedelic experience — including aspects that are known to be predictive of subsequent mental health outcomes. It is important to remark that the present study leveraged prior data to investigate the predictive power of autonomic biomarkers without assessing their causal role. Future work may design new experiments to investigate further their underlying mechanics, and clarify whether the observed autonomic effects are the result of a simultaneous action of psychedelic compounds on the heart and the brain, or if the cardiac effects are secondary to a direct effect on the brain — while assessing potential bidirectional causality between heart and brain activity.

A key ingredient of our analyses was the utilisation of a novel Bayesian method to estimate heart rate dynamics. In contrast to standard frequentist approaches, our method describes heart rate dynamics via a posterior distribution over potential trajectories conditioned on observed inter-beat intervals, which can be effectively sampled. We used these trajectories to estimate Bayesian estimators of the mean HR, HF-HRV, and HR entropy, leading to effective biomarkers. Future work may apply this approach to build Bayesian estimators of other popular metrics, e.g. fractality or other non-linear properties of heart rate dynamics (24, 37), which may be used to explain and track changes linked to a variety of interventions or conditions of interest.

Most studies of the effect of psychedelics on heart activity have focused on sympathetic effects (which are related to actions requiring quick responses), consistently reporting increases in heart rate, systolic blood pressure, and pupil size (45–49). Studies have also focused on the relationship between psychedelic use and cardiac diseases (50, 51). In contrast, the effects of psychedelics on the parasympathetic system (which drives slower autonomic reactions, e.g. relaxation after stress) have been much less investigated. The only systematic investigation of HRV on psychedelics to date — to the best of our knowledge — reported no significant changes in high-frequency HRV (typically associated with parasympathetic activity) after LSD administration (52), suggesting that the acute effects of psychedelics may predominantly rely on sympathetic activity. The differences between previous results and those presented here, particularly related to changes on high-frequency HRV, may be associated with the benefits of the proposed Bayesian approach to estimate HRV.

The observed increase of HR entropy under psychedelics — which captures non-linear aspects of the variability of heart-rate — suggests interesting lines of future investigation related to the parallels between brain and heart entropy. Previous work has proposed HRV as an indicator of health, with this variability being reduced in a wide range of conditions such as hypertension, aortic valve disease, myocardial infarction, diabetes, smoking, and alcohol consumption, to name just a few (23, 53–55). A separate line of investigation has found that increased brain entropy under psychedelics predicts improvements in mental health via an intermediary report of psychological insight (56), suggesting that psychedelics may favour information processing at a greater information granularity. Increased brain entropy under psychedelics may also imply greater system flexibility (31), supporting a learning-based mechanism to psychedelic therapy (19). Furthermore, preliminary evidence indicates that increased entropy may predict subsequent increases in neuronal and psychological plasticity (17, 19, 20). We speculate that these entropic effects in brain and heart activities may reflect a common underlying principle: that diversity in the spontaneous dynamics of living systems offers an adaptive advantage in complex environments (57, 58). Future work with larger sample sizes may investigate dynamical interactions between heart and brain entropies, to elucidate causal (59) or synergistic (60) relationships between. The complementarity of brain and heart entropy to predict certain dimensions of the psychedelic experience is particularly interesting, and future work may explore the mechanisms underlying this phenomenon. Future work may also investigate if the correlated increases between heart and brain entropy are unique to psychedelics, or whether they correspond to broader physiological adjustments. Intriguingly, the correspondence observed between brain entropy and HRV under psychedelics may be disrupted in mental disorders — e.g. brain entropy is increased in schizophrenia (61, 62) and some cases of depression (63, 64) while HRV is reduced (65). Confirming this decoupling between heart and brain entropies and uncovering its driving mechanisms, perhaps using computational approaches similar to Ref. (66), is an interesting line of future work that could deepen our understanding of the relationship between psychedelics and mental health more broadly.

This work represents a first step into exploring the relationship between the altered state of consciousness induced by psychedelics and the autonomic system — a rich space that deserves further investigations. Interestingly, interoceptive feedback has been shown to play a crucial role in the embodied experience and the sense of self (67), as demonstrated in various paradigms where changes in body ownership correspond to changes in the ability to sense heartbeats through bodily sensations (68, 69). Moreover, the role of interoceptive signals in self-related processing has also been demonstrated in neural responses to heartbeats in paradigms related to self-related thoughts (70), mirror self-identification (71), and the embodiment of virtual avatars (72). While analysing neural responses to heartbeats provides valuable insights, the interaction between the brain and the heart also involves highly non-trivial dynamics (73), which have been speculated to be related to ongoing autonomic changes which — in turn — may reflect various dimensions of conscious experience (10). Future work may deepen our understanding of the physiology underlying the altered states of consciousness, e.g. by studying their effect on different markers of the brain-heart interaction, such as heartbeat-evoked potentials (74), phase synchronisation (75), or directed interactions (76). Indeed, it is sometimes argued that peripheral physiological changes observed under psychedelics should be seen as nuisance signal to be regressed from statistical modelling pertaining to acute brain effects; however, an alternative view is that these changes are part of the experience itself, and therefore the corresponding signals may be a bearers of signal rather than noise.

The results presented here have broader implications beyond the psychedelic experience, providing insights on the brain-heart link and its relevance for cognitive neuroscience, psychiatry, and biomedical sciences (10). In particular, the ability to effectively track altered states of consciousness (such as the psychedelic state) from heart signals alone opens new opportunities for a wide range of practical applications. In effect, in contrast to the brain, obtaining reliable and reproducible measures of heart activity is comparatively easier (77), while being less intrusive and significantly more affordable. Future studies may, for example, leverage the proposed techniques to build reliable predictors of specific aspects of experiences under altered states (related to psychedelics or not) that may be used for clinical applications and naturalistic studies. Furthermore, the methods introduced in this work have the potential to provide real-time biofeedback, which could eventually be used by therapists or patients for adjusting the rate of administration of a specific substances — e.g. continuous infusion of DMT (78). Overall, the development of cheap, readily available autonomic biomarkers for mental states has huge practical and clinical potential. We hope that the work presented here will stimulate further efforts to measure and incorporate multi-variate physiological signals beyond the brain, in order to offer a more holistic and comprehensive approach to psychology, psychiatry, and medicine.

## Materials and Methods

### Datasets

The LSD dataset corresponds to previously unpublished data from the experiment originally reported in Refs. (26, 34). Twenty subjects participated in the study by attending two experimental sessions: one in which they received intravenous (i.v.) saline solution (i.e. placebo), and one in which they received i.v. LSD (75 μg). The order of the sessions was randomised, separated by two weeks, and participants were blind to the order (i.e. single blind design). MEG and ECG data were collected under four conditions: resting state with eyes closed, listening with eyes closed to instrumental ambient music (tracks from the album “Eleusian Lullaby” by Alio Die), resting state with eyes open (focusing on a fixation dot), and watching a silent documentary (segments of the “Frozen Planet” documentary series produced by the BBC). LSD was injected approximately four hours prior to the ECG recording.

The DMT dataset corresponds to an experiment first reported in Ref. (33). Thirteen subjects attended two experimental sessions receiving placebo and DMT using a fixed order design, in which subjects received placebo first, followed by DMT. Brain activity was recorded continuously using a 32-channel Brainproducts EEG system, which included an ECG channel. As this was a pilot study, different participants received different doses of DMT (participants were blind to the dose received). Three participants received 7 mg, four received 14 mg, one received 18 mg and five received 20 mg of DMT fumarate in a 2 mL sterile solution over 30 seconds. This was followed by a saline flush lasting 15 seconds. Participants were asked to keep their eyes closed while wearing an eye mask throughout the recordings and low-volume ambient music played in the background to ensure psychological safety.

The psilocybin dataset was first reported in Ref. (79). In this study fifteen healthy male volunteers received placebo and then psilocybin in separate scans conducted on the same day. 275-channel MEG were recorded, including a five minute baseline period and five minutes after infusion of psilocybin. The infusion of psilocybin took approximately 60 seconds and consisted of 2 mg of psilocybin dissolved in 10 mL of saline. Participants were seated in the MEG scanner with their eyes open and asked to fixate on a cross. As this dataset didn’t have a ECG channel, so the cardiac activity was extracted from the MEG data via independent component analysis.

The ketamine dataset was first reported in Ref. (80) and consisted of nineteen healthy male volunteers. On separate days participants received either an infusion of saline or ketamine. The ketamine infusion consisted of an initial bolus of 0.25 mg*/*kg mg/kg delivered over approximately 60 seconds, followed by a maintenance infusion of 0.375 mg/kg/hour. In this experiment, participants lay supine in the MEG scanner while 275-channel MEG and ECG was recorded with their eyes open and fixating on a cross. Five minutes of baseline was recorded prior to the infusion commencing, and ten minutes from the commencement of the infusion.

### Data preprocessing

Inter-beat intervals were estimated from the ECG via a semi-automated procedure where first the ECG was processed via a wavelet transform, and then peaks where detected using the `findpeaks` function in MATLAB (v R2020b) with manually adjusted minimum height and minimum inter-beat distance. Then, an ectopic correction was performed, where a heart-beat interval was declared abnormal if its length was more than 20% different from the previous one. Datasets with more than 15% of ectopic beats were rejected for being too noisy — leading to the rejection of only 4 time series from the ketamine dataset (four different subjects, two in drug and two in placebo conditions). Otherwise, ectopic beats were corrected by replacing them for the linear interpolation between the previous and successive beat. The resulting sequence of inter-beat intervals was used to estimate heart rate dynamics, either via the Bayesian or the frequentist approach.

### Bayesian estimation of heart rate dynamics

Here we provide an overview of our approach to estimate heart rate dynamics — full technical details can be found in Ref. (81).

The conventional, frequentist method to calculate instantaneous heart rate is to estimate the inter-beat intervals *I*_b_, and then compute HR_freq_(*t*) = 60*/I*_b_(*t*). From a statistical perspective, this expression can be understood as the outcome of an elementary method of inference that delivers as estimate the number of beats one would see if all beats were separated by the same inter-beat interval *I*_b_. In contrast, our Bayesian framework conceives the heart rate as a hidden process that drives the actual observed heartbeats, the statistical properties of which can be estimated via generative modelling. Our modelling approach is based on two time series that correspond to the values of dynamical processes with sampling frequency *f*_s_ =1 Hz: *x*_*t*_, which counts the number of heart beats in a temporal bin of length ∆*t* (i.e. between the present moment and the previous sample), and *z*_*t*_, which represents the heart rate that drives the corresponding heart beats (see Figure 5). In general, we assume that the heart beats *x*_*t*_ are observed, and that sequence of heart rates *z*_*t*_ follows a hidden stochastic process whose statistical properties can nevertheless be inferred.

**Fig. 5.**
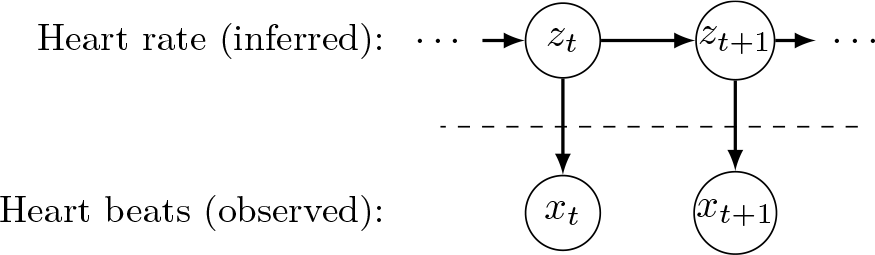
Proposed heart rate state-space modelling approach. The observable data (the heart beats, *x*_*t*_) are assumed to be driven by the dynamics of a hidden stochastic process (the heart rate, *z*_*t*_), which cannot be directly measured but can be inferred from the data.

**Fig. 6.**
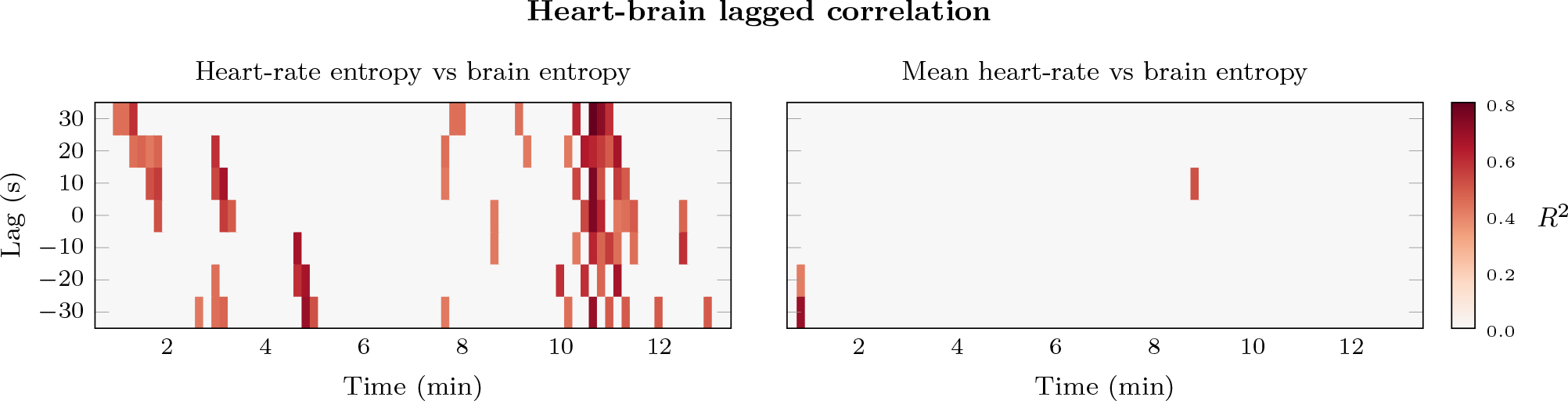
Thresholded results of predictive power of heart-to-brain biomarker. Results as in Figure 3a, but here the *R*^2^ values are thresholded using non-parametric confidence intervals. Specifically, *R*^2^ values were calculated for each timepoint, but shuffling the values of heart-rate and brain pairs of data points. Shuffling is done 1000 times per timepoint, which is used to estimate the resulting values of *R*^2^ under a null distribution. Finally, the 95% quantile is of this distribution is calculated, which is used to threshold the original *R*^2^ values.

In order to build a joint probability distribution over heart beats *x*_1_, …, *x*_*T*_ and heart rates *z*_1_, …, *z*_*T*_, we follow the state-space literature (82) in adopting the following assumptions: (i) the dynamics of the heart rate are Markovian; and (ii) given the value of heart rate *z*_*t*_, the number of heartbeats at that given temporal bin *x*_*t*_ is conditionally independent of other heart rate values at different timepoints. These assumptions allow us to express the joint distribution of heart rate and heartbeat counts as:

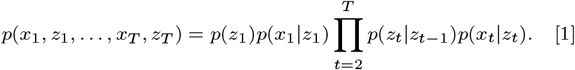

Therefore, the specification of the full model requires only three ingredients: the heart rate dynamics in the form of the conditional probability *p*(*z*_*t*_ |*z*_*t*−1_), the link between heart rate and heart beats *p*(*x*_*t*_ |*z*_*t*_), and the distribution of the initial condition *p*(*z*_1_). As described in Ref. (81), we model *p*(*x*_*t*_|*z*_*t*_) using a Poisson distribution and a small bin size in order to guarantee good statistical fit, while we model *p*(*z*_*t*_ |*z*_*t* −1_) and *p*(*z*_1_) using a Gamma Markov chain (GMC) (83, 84). Using the resulting generative model *p*(*x*_1_, *z*_1_, …, *x*_*T*_, *z*_*T*_) and Bayes’ rule, one can describe heart rate dynamics not via a point estimate (i.e. as a single, most likely trajectory) but as following the posterior distribution:

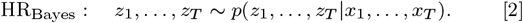

This posterior distribution describes the most likely heart rate trajectories *z*_1_, …, *z*_*N*_ given the observed data *x*_1_, …, *x*_*N*_.

Thanks to these modelling choices, *p*(*z*_1_, …, *z*_*T*_| *x*_1_, …, *x*_*T*_) can be efficiently sampled via a Gibbs sampler (85) — proposed in Ref. (86). Our experiments suggest that the resulting model is generally non-ergodic, which motivated the following sampling procedure. To generate one trajectory, we run the Gibbs sampler *N*_r_ = 2 ×10^4^ iterations, and discard the first *N*_d_ = 5 ×10^3^. We calculate the mean value of the remaining *N*_r_ − *N*_d_ runs and take the result as a single trajectory. For each sequence of heartbeats (corresponding to different drugs, subjects, and experimental conditions), we iterate this procedure *N*_s_ = 500 times, initialising the Gibbs sampler every time with new random initial conditions. As discussed in Ref. (81), the Gibbs sampler depends on two hyper-parameters *θ* and *τ* that determine the prior and estimation of a connectivity strength parameter of the Gamma Markov chain denoted by *γ*. For all datasets we use *θ* = 10 and *τ* = 1. The sampling of heart rate trajectories was done using a combination of Python (v. 3.9.1) and Julia (v. 1.5.3), using code available in the repository github.com/ferosas/BayesianAtHeart.

### Bayesian estimators of properties of heart dynamics

We used the sampled trajectories of heart rate dynamics to build Bayesian estimators of properties of heart rate dynamics. To explain this part of the method, we introduce the shorthand notation ***z*** = (*z*_1_, …, *z*_*T*_) and ***x*** = (*x*_1_, …, *x*_*T*_) for sampled trajectories of heart rate and heart beats respectively, and let *F* (***z***) be a scalar function of this trajectory — i.e. any scalar property of the heart rate trajectory, such as its mean value or entropy. Then, the generative model above can be used to derive the posterior distribution of the property *F*, which corresponds to *p*(*F* (***z***) | ***x***). Sampled trajectories can be used to estimate various properties of this posterior — e.g. its mean:

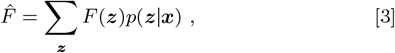

where the value of the property *F* for each possible trajectory is weighted by the likelihood of such trajectory given the observed data. If *F* is a linear property, then Eq. (3) accepts a shortcut: it reduces to 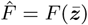, where 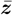 is the average trajectory under the posterior *p*(***z***|***x***). We used this shortcut to build Bayesian estimators of mean heart rate and its low and high frequency components. To extract the power in low and high-frequencies in the average spectrum of the HR dynamics, we calculated a spectral decomposition of the variance of the average HR trajectory. Specifically, we first filtered the average heart rate trajectories using a bandpass Butterworth filter of order 2 over standard bands (0.04–0.15 Hz for low-frequency and 0.15–4 Hz for high-frequency components), and then calculated the variance of the resulting signal. The results in the first three columns in Figure 2 use HRV calculated over partially overlapping sliding windows of 60 seconds separated by 1 second.

### Estimating heart rate entropy

Brain entropy in M/EEG data is usually calculated via Lempel-Ziv complexity (referred to as LZ), which estimates how diverse the patterns exhibited by a given signal are (30, 34). The method was introduced by engineers Abraham Lempel and Jacob Ziv to study the statistics of binary sequences (39) (later becoming the basis of the well-known`zip` compression algorithm), and has been used to study patterns in M/EEG activity for more than 20 years. It has become a popular method to track states of consciousness, displaying consistent decreases under anaesthesia (87, 88) and sleep (89), and increases during psychedelics (32, 33), meditation (90),^†^ and states of ‘flow’ associated with music improvisation (92).

There are two issues that make it challenging to apply LZ complexity to heart rate data: the need of a binarisation step combined with the non-stationarity of the time series (observed here in all drugs except LSD), and the relatively short length of the time series (LZ is usually estimated on M/EEG data over windows of thousands of samples, which is challenging with a sampling frequency of 1 Hz). We addressed these challenges with two innovations, which we explain the following.

First, entropy estimation in M/EEG requires the binarisation of the data, which is often done by thresholding on the signal’s mean value. While this particular choice usually doesn’t have a big impact on the entropy estimates (87), it becomes problematic with highly non-stationary data, as it could lead to an underestimation of the entropy due to long periods of the signal being either above or bellow its mean value. To avoid this problem, instead of thresholding based on the mean value, we threshold according to the sign of the derivative — hence a ‘1’ implies the signal is increasing and a ‘0’ that it is decreasing.

To address the issue of the low sampling frequency and resulting short length of the time series data, we don’t use the classic LZ algorithm (93) but instead estimate entropy using the *Context Tree Weighted* (CTW) algorithm (38). This algorithm has shown to converge quicker than other entropy rate estimators including LZ (94). Crucially, the flexibility of the CTW algorithm allows it to be trained and tested with different datasets: the training dataset is used to estimate the likelihood of different patterns, while the test data is used to calculate their frequency. We use this powerful feature to train the algorithm on all the observed patterns during the whole placebo session, and test it during small windows during either drug or placebo sessions. Results reported in Figure 2 were obtained using windows of 120 samples. The CTW algorithm uses one free parameter, which determine the maximum length of patterns that it accounts for; for simplicity, we set that length to be equal to the window length.

### Calculating brain entropy

For calculating the brain entropy on the DMT EEG data, we used the CTW algorithm to train and test over partially overlapping sliding windows of 5 seconds separated from each other by 1 second. The EEG data was sampled at 1000 Hz, which resulted in 5000 samples per window. This procedure yielded one entropy estimate per electrode per second, with the values of entropy during discarded data being imputed via linear interpolation over time. The resulting values were averaged over electrodes, giving a single time series of entropy values per subject with one sample per second.

For the LSD MEG data, we computed source-reconstructed signals on the centroids of the AAL-90 atlas (42), following the procedure in Ref. (34). Data was sampled at 600 Hz and divided in 2 s epochs, resulting in windows with 1200 samples. We performed mean binarisation, computed CTW for each epoch, subject, and AAL region, and finally averaged across epochs to obtain one CTW value per subject and region.

### Statistical analysis

Cluster statistics (41) for the results in Figure 2 were calculated as follows. Each time series of heart rate properties (mean HR, low-frequency HRV, high-frequency HRV, heart entropy) was first normalised by subtracting its baseline value — defined as the mean value obtained 100 seconds prior to the drug injection. Then, for each drug (DMT, psilocybin, and ketamine) and each condition (drug or placebo), a one-sample t-test was performed; then an adaptive threshold (41) was used to determine cluster candidates; then a cluster statistic (sum of t-scores of cluster members) was calculated per cluster. Finally, the significance of each cluster was determined by repeating the same process 500 times randomly flipping the sign of the values, and taking the maximal cluster statistic to build a null distribution against which the original cluster scores were contrasted against.

The explained variance calculated in Figure 2 corresponds to the *R*^2^ of a rank regression model. Note that this is equivalent to the square of the Spearmann correlation, and hence is robust to outliers.

Linear mixed models were used to estimate the relationship between HR and brain entropy. For this, we constructed a model using HR entropy as dependent variable, brain entropy (z-scored), condition (drug vs placebo) and their interaction as fixed effects, and added non-nested random intercepts for subject ID and setting (eyes closed, eyes open, music, and video). In standard Wilkinson notation,

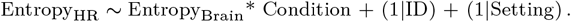

The significance of this model was estimated by calculating a log-likelihood ratio test against a model that is identical but does not include brain entropy as fixed effect. The resulting p-value was FDR-corrected via a standard Benjamini-Hochberg procedure (43). Figure 4 presents the value of the estimates of the interaction between brain entropy and condition, which reflects the effect of brain entropy changes over HR entropy during the drug condition.

Linear mixed-effects modelling was also used to quantify the predictive power of brain and autonomic markers over subjective scores. When considering individual markers we built models using a given visual analog score (VAS) as dependent variable, the marker, condition, and their interaction as fixed effects, and non-nested random intercepts for subject ID and setting. As before, the estimate reported in Figure 4 is the interaction between the biomarker and condition. When considering two markers, the model was the same but considered the two markers, condition, and their triple interaction as fixed effect, with the estimate of the triple interaction being reported in Figure 4.

All linear mixed models were implemented in R Studio (v. 2022.12.0+353) using the packages `lmer` (95) and `lmerTest` (96).

## ACKNOWLEDGMENTS

F.R. is supported by the Fellowship Programme of the Institute of Cultural and Creative Industries of the University of Kent. A.I.L. is supported by the Molson Neuro-Engineering Fellowship. D.B. is funded by the Wellcome Trust (grant no. 646 210920/Z/18/Z).

## Supplementary Materials

This section contains Supplementary Figure 6, supporting the results on the predictive power of hear-brain biomarkers.

## Footnotes

∗ We didn’t include LF-HRV, as it didn’t exhibit consistent changes in the previous analyses.

† Note that in meditation the effect depends on years of experience, see Ref. (91) for a review.

## References

1. FJ Varela, E Thompson, E Rosch, J Kabat-Zinn, The Embodied Mind: Cognitive Science and Human Experience. (MIT press), (2017).

2. AK Seth, Interoceptive inference, emotion, and the embodied self. Trends Cogn. Sci. 17, 565–573 (2013).

3. M Tsakiris, H De Preester, The Interoceptive Mind: From Homeostasis to Awareness. (Oxford University Press), (2018).

4. JF Thayer, E Sternberg, Beyond heart rate variability. Annals New York Acad. Sci. 1088, 361–372 (2006).

5. GG Berntson, PJ Gianaros, M Tsakiris, Interoception and the autonomic nervous system: Bottom-up meets top-down. (Oxford University Press), (2019).

6. HD Critchley, NA Harrison, Visceral influences on brain and behavior. Neuron 77, 624–638 (2013).

7. HD Critchley, SN Garfinkel, Interoception and emotion. Curr. Opin. Psychol. 17, 7–14 (2017).

8. AD Craig, How do you feel – now? The anterior insula and human awareness. Nat. Rev. Neurosci. 10, 59–70 (2009).

9. M Tsakiris, My body in the brain: A neurocognitive model of body-ownership. Neuropsychologia 48, 703–712 (2010).

10. D Candia-Rivera, Brain-heart interactions in the neurobiology of consciousness. Curr. Res. Neurobiol. 3, 100050 (2022).

11. FX Vollenweider, KH Preller, Psychedelic drugs: Neurobiology and potential for treatment of psychiatric disorders. Nat. Rev. Neurosci. 21, 611–624 (2020).

12. C Timmermann, et al., A neurophenomenological approach to non-ordinary states of consciousness: Hypnosis, meditation, and psychedelics. Trends Cogn. Sci. (2022).

13. R Cofré, et al., Whole-brain models to explore altered states of consciousness from the bottom up. Brain Sci. 10, 626 (2020).

14. MW Johnson, PS Hendricks, FS Barrett, RR Griffiths, Classic psychedelics: An integrative review of epidemiology, therapeutics, mystical experience, and brain network function. Pharmacol. & Ther. 197, 83–102 (2019).

15. R Kočárová, J Horáček, R Carhart-Harris, Does psychedelic therapy have a transdiagnostic action and prophylactic potential? Front. Psychiatry p. 1068 (2021).

16. RL Carhart-Harris, et al., The entropic brain: A theory of conscious states informed by neuroimaging research with psychedelic drugs. Front. Hum. Neurosci. p. 20 (2014).

17. RL Carhart-Harris, KJ Friston, REBUS and the anarchic brain: Toward a unified model of the brain action of psychedelics. Pharmacol. Rev. 71, 316–344 (2019).

18. LR Swanson, Unifying theories of psychedelic drug effects. Front. Pharmacol. p. 172 (2018).

19. R Carhart-Harris, et al., Canalization and plasticity in psychopathology. Neuropharmacology 226, 109398 (2022).

20. I Hipólito, J Mago, F Rosas, R Carhart-Harris, Pattern breaking: A complex systems approach to psychedelic medicine. PsyArXiv (2022).

21. C Letheby, Philosophy of psychedelics. (Oxford University Press), (2021).

22. GD Clifford, Ph.D. thesis (Oxford University, UK) (2002).

23. UR Acharya, K Paul Joseph, N Kannathal, CM Lim, JS Suri, Heart rate variability: A review. Med. & Biol. Eng. & Comput. 44, 1031–1051 (2006).

24. G Valenza, L Citi, JP Saul, R Barbieri, Measures of sympathetic and parasympathetic autonomic outflow from heartbeat dynamics. J. Appl. Physiol. 125, 19–39 (2018).

25. FX Vollenweider, M Kometer, The neurobiology of psychedelic drugs: Implications for the treatment of mood disorders. Nat. Rev. Neurosci. 11, 642–651 (2010).

26. RL Carhart-Harris, et al., Neural correlates of the LSD experience revealed by multimodal neuroimaging. Proc. Natl. Acad. Sci. 113, 4853–4858 (2016).

27. R McMillan, SD Muthukumaraswamy, The neurophysiology of ketamine: An integrative review. Rev. Neurosci. 31, 457–503 (2020).

28. C Timmermann, et al., Human brain effects of DMT assessed via EEG-fMRI. Proc. Natl. Acad. Sci. 120, e2218949120 (2023).

29. E Drummond, et al., Psychedelic resting-state neuroimaging: A review and perspective on balancing replication and novel analyses. Neurosci. & Biobehav. Rev. 138, 104689 (2022).

30. RL Carhart-Harris, The entropic brain – revisited. Neuropharmacology 142, 167–178 (2018).

31. M Girn, et al., A complex systems perspective on psychedelic brain action. Trends Cogn. Sci. (2023).

32. MM Schartner, RL Carhart-Harris, AB Barrett, AK Seth, SD Muthukumaraswamy, Increased spontaneous MEG signal diversity for psychoactive doses of ketamine, LSD and psilocybin. Sci. reports 7, 46421 (2017).

33. C Timmermann, et al., Neural correlates of the DMT experience assessed with multivariate EEG. Sci. Reports 9, 1–13 (2019).

34. PA Mediano, et al., Effects of external stimulation on psychedelic state neurodynamics. BioRxiv pp. 2020–11 (2020).

35. F Beckers, B Verheyden, AE Aubert, Aging and nonlinear heart rate control in a healthy population. Am. J. Physiol. Circ. Physiol. 290, H2560–H2570 (2006).

36. R Acharya U, OW Sing, LY Ping, T Chua, Heart rate analysis in normal subjects of various age groups. Biomed. Eng. Online 3, 1–8 (2004).

37. F Shaffer, JP Ginsberg, An overview of heart rate variability metrics and norms. Front. Public Heal. p. 258 (2017).

38. FM Willems, YM Shtarkov, TJ Tjalkens, The context-tree weighting method: Basic properties. IEEE Transactions on Inf. Theory 41, 653–664 (1995).

39. A Lempel, J Ziv, On the complexity of finite sequences. IEEE Transactions on Inf. Theory 22, 75–81 (1976).

40. PA Mediano, et al., Spectrally and temporally resolved estimation of neural signal diversity. bioRxiv pp. 2023–03 (2023).

41. E Maris, R Oostenveld, Nonparametric statistical testing of EEG- and MEG-data. J. Neurosci. Methods 164, 177–190 (2007).

42. N Tzourio-Mazoyer, et al., Automated anatomical labeling of activations in SPM using a macroscopic anatomical parcellation of the MNI MRI single-subject brain. NeuroImage 15, 273–89 (2002).

43. Y Benjamini, Y Hochberg, Controlling the false discovery rate: A practical and powerful approach to multiple testing. J. Royal Stat. Soc. Ser. B (Methodological) 57, 289–300 (1995).

44. SD Kreibig, Autonomic nervous system activity in emotion: A review. Biol. psychology 84, 394–421 (2010).

45. HJ Deshon, M Rinkel, HC Solomon, Mental changes experimentally produced by LSD (d-lysergic acid diethylamide tartrate). Psychiatr. Q. (1952).

46. M Kaelen, et al., LSD enhances the emotional response to music. Psychopharmacology 232, 3607–3614 (2015).

47. Y Schmid, et al., Acute effects of lysergic acid diethylamide in healthy subjects. Biol. Psychiatry 78, 544–553 (2015).

48. PC Dolder, Y Schmid, M Haschke, KM Rentsch, ME Liechti, Pharmacokinetics and concentration-effect relationship of oral LSD in humans. Int. J. Neuropsychopharmacol. 19, pyv072 (2016).

49. F Holze, et al., Distinct acute effects of LSD, MDMA, and D-amphetamine in healthy subjects. Neuropsychopharmacology 45, 462–471 (2020).

50. O Simonsson, W Osika, R Carhart-Harris, PS Hendricks, Associations between lifetime classic psychedelic use and cardiometabolic diseases. Sci. reports 11, 14427 (2021).

51. LM Salinsky, et al., μ-opioid receptor agonists and psychedelics: Pharmacological opportunities and challenges. Front. Pharmacol. 14, 1239159 (2023).

52. S Olbrich, KH Preller, FX Vollenweider, LSD and ketanserin and their impact on the human autonomic nervous system. Psychophysiology 58, e13822 (2021).

53. MA Woo, WG Stevenson, DK Moser, HR Middlekauff, Complex heart rate variability and serum norepinephrine levels in patients with advanced heart failure. J. Am. Coll. Cardiol. 23, 565–569 (1994).

54. PK Stein, PP Domitrovich, N Hui, P Rautaharju, J Gottdiener, Sometimes higher heart rate variability is not better heart rate variability: Results of graphical and nonlinear analyses. J. Cardiovasc. Electrophysiol. 16, 954–959 (2005).

55. M Valentini, G Parati, Variables influencing heart rate. Prog. Cardiovasc. Dis. 52, 11–19 (2009).

56. T Lyons, et al., Enduring human brain changes after psilocybin. under revision (2023).

57. JP Crutchfield, Between order and chaos. Nat. Phys. 8, 17–24 (2012).

58. A Brouwer, RL Carhart-Harris, Pivotal mental states. J. Psychopharmacol. 35, 319–352 (2021).

59. D Janzing, D Balduzzi, M Grosse-Wentrup, B Schölkopf, Quantifying causal influences. PsyArXiv (2013).

60. PA Mediano, F Rosas, RL Carhart-Harris, AK Seth, AB Barrett, Beyond integrated information: A taxonomy of information dynamics phenomena. arXiv preprint arXiv:1909.02297 (2019).

61. A Fernández, et al., Lempel–ziv complexity in schizophrenia: A MEG study. Clin. Neurophysiol. 122, 2227–2235 (2011).

62. Y Li, et al., Abnormal EEG complexity in patients with schizophrenia and depression. Clin. Neurophysiol. 119, 1232–1241 (2008).

63. M Bachmann, K Kalev, A Suhhova, J Lass, H Hinrikus, Lempel Ziv complexity of EEG in depression in 6th European Conference of the International Federation for Medical and Biological Engineering: MBEC 2014, 7-11 September 2014, Dubrovnik, Croatia. (Springer), pp. 58–61 (2015).

64. M. Méndez, et al., Complexity analysis of spontaneous brain activity: Effects of depression and antidepressant treatment. J. psychopharmacology 26, 636–643 (2012).

65. JM Montaquila, BJ Trachik, JS Bedwell, Heart rate variability and vagal tone in schizophrenia: A review. J. Psychiatr. Res. 69, 57–66 (2015).

66. H Rajpal, et al., Psychedelics and schizophrenia: Distinct alterations to bayesian inference. NeuroImage 263, 119624 (2022).

67. K Suzuki, SN Garfinkel, HD Critchley, AK Seth, Multisensory integration across exteroceptive and interoceptive domains modulates self-experience in the rubber-hand illusion. Neuropsychologia 51, 2909–2917 (2013).

68. V Ainley, A Tajadura-Jiménez, A Fotopoulou, M Tsakiris, Looking into myself: Changes in interoceptive sensitivity during mirror self-observation. Psychophysiology 49, 1672–1676 (2012).

69. ML Filippetti, M Tsakiris, Heartfelt embodiment: Changes in body-ownership and self-identification produce distinct changes in interoceptive accuracy. Cognition 159, 1–10 (2017).

70. M Babo-Rebelo, CG Richter, C Tallon-Baudry, Neural responses to heartbeats in the default network encode the self in spontaneous thoughts. J. Neurosci. 36, 7829–7840 (2016).

71. A Sel, RT Azevedo, M Tsakiris, Heartfelt self: Cardio-visual integration affects self-face recognition and interoceptive cortical processing. Cereb. Cortex 27, 5144–5155 (2017) Publisher: Oxford Academic.

72. HD Park, et al., Transient modulations of neural responses to heartbeats covary with bodily self-consciousness. J. Neurosci. 36, 8453–8460 (2016).

73. K Schiecke, et al., Brain-heart interactions considering complex physiological data: Processing schemes for time-variant, frequency-dependent, topographical and statistical examination of directed interactions by convergent cross mapping. Physiol. Meas. 40, 114001 (2019).

74. HD Park, O Blanke, Heartbeat-evoked cortical responses: Underlying mechanisms, functional roles, and methodological considerations. NeuroImage 197, 502–511 (2019).

75. M Dumont, et al., Interdependency between heart rate variability and sleep EEG: Linear/non-linear? Clin. Neurophysiol. 115, 2031–2040 (2004).

76. D Candia-Rivera, Modeling brain-heart interactions from Poincaré plot-derived measures of sympathetic-vagal activity. MethodsX 10, 102116 (2023).

77. RE Kleiger, et al., Stability over time of variables measuring heart rate variability in normal subjects. The Am. J. Cardiol. 68, 626–630 (1991).

78. L Luan, et al., Psychological and physiological effects of extended DMT. PsyArXiv (2023).

79. SD Muthukumaraswamy, et al., Broadband cortical desynchronization underlies the human psychedelic state. J. Neurosci. 33, 15171–15183 (2013).

80. SD Muthukumaraswamy, et al., Evidence that subanesthetic doses of ketamine cause sustained disruptions of NMDA and AMPA-mediated frontoparietal connectivity in humans. J. Neurosci. 35, 11694–11706 (2015).

81. FE Rosas, D Candia-Rivera, AI Luppi, Y Guo, PA Mediano, Bayesian at heart: Towards autonomic outflow estimation via generative state-space modelling of heart rate dynamics. arXiv preprint arXiv:2303.04863 (2023).

82. J Durbin, SJ Koopman, Time Series Analysis by State Space Methods. (Oxford University Press) Vol. 38, (2012).

83. AT Cemgil, O Dikmen, Conjugate gamma Markov random fields for modelling nonstationary sources in International Conference on Independent Component Analysis and Signal Separation. (Springer), pp. 697–705 (2007).

84. O Dikmen, AT Cemgil, Unsupervised single-channel source separation using Bayesian NMF in 2009 IEEE Workshop on Applications of Signal Processing to Audio and Acoustics. (IEEE), pp. 93–96 (2009).

85. CM Bishop, Pattern Recognition and Machine Learning. (Springer), (2006).

86. S Gugushvili, F van der Meulen, M Schauer, P Spreij, Fast and scalable non-parametric Bayesian inference for poisson point processes. arXiv preprint arXiv:1804.03616 (2018).

87. XS Zhang, RJ Roy, EW Jensen, EEG complexity as a measure of depth of anesthesia for patients. IEEE Transactions on Biomed. Eng. 48, 1424–1433 (2001).

88. M Schartner, et al., Complexity of multi-dimensional spontaneous EEG decreases during propofol induced general anaesthesia. PloS ONE 10, e0133532 (2015).

89. MM Schartner, et al., Global and local complexity of intracranial EEG decreases during NREM sleep. Neurosci. Conscious. 2017, niw022 (2017).

90. RM Vivot, C Pallavicini, F Zamberlan, D Vigo, E Tagliazucchi, Meditation increases the entropy of brain oscillatory activity. Neuroscience 431, 40–51 (2020).

91. DA Atad, PA Mediano, F Rosas, A Berkovich-Ohana, Meditation and complexity: A systematic review. PsyArXiv (2023).

92. D Dolan, et al., The improvisational state of mind: A multidisciplinary study of an improvisatory approach to classical music repertoire performance. Front. Psychol. 9, 1341 (2018).

93. F Kaspar, H Schuster, Easily calculable measure for the complexity of spatiotemporal patterns. Phys. Rev. A 36, 842 (1987).

94. Y Gao, I Kontoyiannis, E Bienenstock, Estimating the entropy of binary time series: Methodology, some theory and a simulation study. Entropy 10, 71–99 (2008).

95. D Bates, M Mächler, B Bolker, S Walker, Fitting linear mixed-effects models using lme4. J. Stat. Softw. 67, 1–48 (2015).

96. A Kuznetsova, PB Brockhoff, RHB Christensen, lmerTest package: Tests in linear mixed effects models. J. Stat. Softw. 82, 1–26 (2017).

